# Morphometric and elastic properties of immature reticulocytes in health and during acute lymphoblastic and acute myeloid leukemia

**DOI:** 10.1101/2020.11.05.369868

**Authors:** Evgeniy S. Seliverstov

## Abstract

Despite significant advances, many changes occurring in the tumor microenvironment during acute lymphoblastic leukemia (ALL) and acute myeloid leukemia (AML) remain unclear. The surface of immature reticulocytes was examined by atomic force microscopy (AFM) to determine specific changes during the development of ALL and AML. In patients with ALL the surface area of reticulocytes increased by 18.5%, volume by 8.7%, the width of invaginations by 18%, and cell height decreased by 7.8%. In patients with AML, the volume increased by 12.6%, roughness by 35.5%, the height of protrusions by 36.2%, the depth of invaginations by 24.8%, their width by 18.2%, and the maximum height difference of the surface by 31.9%. No significant differences in Young’s modulus were found, but a downward trend was noted. The obtained data has important prognostic value in studying the bone marrow activity during acute leukemia.

**Highlights:** AFM reveals implicit signs of malignant disease development in the blood. ALL increases area, volume, and invaginations, but reduces the height of IRF. AML increases volume, roughness, protrusions, and invaginations of IRF.

## Introduction

Acute lymphoblastic (or lymphocytic) leukemia (ALL) and acute myeloid leukemia (AML) are the group of blood cell cancers characterized by the appearance of leukemia cells (blasts) — immature lymphocytes (in case of ALL) and immature hematopoietic cells (in case of AML) in the bone marrow and blood. These abnormal cells affect normal blood cell precursors and gradually inhibit their growth and maturation^1-2^. ALL more commonly occurs in those 15 to 39 years of age, while AML more frequently occurs among people aged 65-74. In recent years, rates for new cases of ALL adjusted by age have been rising on average by 0.6% each year while death rates have been decreasing on average by 1.0% each year with 68.8% 5-year relative survival rate in the United States^3^. Rates for new cases of AML have been also rising with an average increase of 1.5% each year, but death rates remain stable over 2008-2017 with 28.7% of 5-year relative survival rate^4^.

Very few epidemiological studies of acute leukemia and long-term monitoring of the vital status of patients have been carried out in Russia. Recent researches provide that according to estimates, the 3-year survival rate of ALL patients in Russia is 47% and the 3-year survival rate of AML patients is only 20%^5^. These data show that despite significant advances in the early diagnosis and treatment of ALL and AML, we still need a deeper understanding of the malignant processes that occur during their development, and the compensatory-adaptive processes of human body that arise in response to them.

Leukemic processes usually begin in the bone marrow causing inhibition of normal erythropoiesis. Therefore, both in the case of studying the earliest changes that occur due to the influence of blasts on their environment and in predicting the course of a disease and success of treatment, it is necessary to have a tool which could be an early indicator of the bone marrow activity.

Reticulocytes are the last precursors of erythrocytes. They are larger in size and volume and have irregular shape. They contain a filamentous-reticular substance (reticulum) comprising residual RNA and components of degraded organelles. Four maturing stages of reticulocytes can be distinguished with the rough estimation of stained reticulum morphology: R1 — reticulum is= densely clumped, R2 — reticulum is presented as a loose and extended network on the whole area of a cell, R3 — reticulum is presented as a residual network with scattered granules, R4 — cells with reticulum are presented as individual strands or scattered granules^6-7^. We can also sort populations of reticulocytes into four sequential subpopulations by flow cytometry based on transferrin receptor (CD71) expression or into three sequential subpopulations based on RNA fluorescence intensity^8-9^.

Immature (R1 and R2) groups of reticulocytes circulate in the peripheral blood flow of healthy individuals in small quantities. But in periods of increased erythropoietic need, the immature reticulocyte fraction (IRF) is released into the bloodstream leading to rapid replenishment of the red blood cell (RBC) population. The IRF major clinical applications as a very sensitive parameter include measuring early hematopoietic recovery after intensive chemotherapy and an indication of the adequacy of response to erythropoietin therapy in patients with anemia that often associated with acute leukemia^10-11^. Therefore, the characterization of specific properties of reticulocytes seems to be a promising research subject as they could help predict therapeutic response to therapies and a better understanding of malignant processes in blood^12^.

The development and implementation of atomic force microscopy (AFM) method in biological researches opened opportunities for studying the properties of native and fixed cells at a qualitatively new level. In particular, researchers can get a high-resolution 3D image of a sample surface and analyze its local properties such as Young’s modulus, roughness, adhesive forces, etc^13-14^. AFM allows researchers to assess the dynamics of rearrangement of membrane structures and reveal the implicit parameters of norm and pathology, which have great importance in studying blood cells that are involved in many compensatory-adaptive processes of the body^15^.

While searching for studies similar to this research, where AFM is used to examine reticulocytes, the only articles found were devoted to studying the life cycle of *Plasmodium* species that infect the reticulocytes population. For instance, authors used AFM to illustrate the presence and density of clathrin pits on uninfected reticulocytes and caveolae in cells with the *P. vivax*^7^. And only one research was found where AFM was used to noninfected reticulocytes to quantitatively estimate shrinkage of spectrin filaments during reticulocyte maturation^16^.

In the present paper, I used different AFM techniques to evaluate the influence of the ALL and AML development on the morphometric and elastic properties of the surface of immature reticulocytes.

The major goals of this research are: (i) to evaluate the elastic properties of reticulocytes by measuring Young’s modulus of cells; (ii) to reveal the morphometric features of reticulocytes by measuring their height, surface area, and volume; (iii) to analyze the surface relief of reticulocytes by measuring the parameters of cell surface roughness; (iv) to study membrane surface structures of reticulocytes by counting the number of protrusions and invaginations on the cell’s membrane surface, and measuring the height of protrusions and depth and width of invaginations.

## Materials and methods

### 2.1 Sample preparation

The study was approved by the local Ethical Committee of the Medical Institute of Belgorod State National Research University and informed consent of all subjects was obtained according to the recommendations of the Helsinki Declaration (The International Response to Helsinki VI, The WMA’s Declaration of Helsinki on Ethical Principles for Medical Research Involving Human Subjects, as adopted by the 52nd WMA General Assembly, Edinburg, October 2000).

Samples of venous blood were taken from patients of both sexes with ALL (N = 10) and AML (N = 10) undergoing medical treatment and healthy donors (N = 10) underwent a clinical examination in the hematological department of the Belgorod regional clinical hospital. Patients and donors both were having an age of 25–45 years on the moment of sampling. Blood sampling was performed by venipuncture by specialized medical staff. Samples were collected in Vacuette K3E vacuum tubes containing dry EDTA K3 at a concentration of 2.0 mg (0.006843 mol/L) per 1 mL of blood.

Immediately on the day of sampling reticulocytes were stained with 0.1% solution of brilliant cresyl blue prepared in 0.9% NaCl solution. The smears were made by the manual spreading of the carefully shaken blood mixed with the dye in a 1:1 ratio by volume. After that, smears were kept unfixed and air-dried.

### 2.2 Atomic force microscopy

The AFM measurements of the surfaces of the reticulocytes were performed in the air, at room temperature immediately after the smear’s drying. Each sample measurement lasted for approximately 3-4 h. AFM imaging was performed using NTEGRA Vita atomic force microscope (configuration based on Olympus IX-71 inverted optical microscope; NT-MDT, Russia). The scanning process was carried out in the contact mode and in the contact error mode with a scanning frequency of 0.8 Hz using gold-plated probes of CSG11 series (NT-MDT, Russia) with a stiffness of 1.1 N/m, the radius of curvature of 10 nm, and nominal spring constant of 5.546 N/m (appendix fig. 1). All images were processed in Nova software (version 1.0.26.1443, NT-MDT, Russia).

Scanning took several steps. First, using an optical microscope, reticulocytes of R1 or R2 maturity classes according to Heilmeyer^6^ was found on the smear (appendix fig. 2). Then a cell was selected and scanned within a 30 μm^2^ area containing it. After that, the size of the scanned area was adjusted to the size of the selected cell. Then, in the center of the cell, a 3×3 μm^2^ area was scanned. And in the end, Young’s modulus of the cell was measured. The images were flattened and plane fitted before analysis.

At least 10 cells were scanned from each sample. Thus, the total number of scanned cells was: 100 in the group of AML patients, 100 in the group of ALL patients and 100 in the group of healthy donors.

After excluding the samples with damage or possible alteration because of the measurement procedure, the final number of collected data was: 93 cells in the group of AML patients, 93 cells in the group of ALL patients and 78 cells in the group of healthy donors.

### 2.3 Young’s modulus

To determine the elastic properties of the reticulocytes, the method of atomic force spectroscopy was used. To measure the Young’s modulus, the cells were scanned in the mode of force spectroscopy when a load was applied to 25 local points of the cell surface (appendix fig. 7). Force curves were obtained and used to calculate the force of interaction between the probe and the sample. The arithmetic mean of Young’s modulus was calculated for each cell. Young’s modulus of the sample-probe system was calculated based on Capella’s formula^17^.

### 2.4 Morphometric properties

The volume (V, μm^3^) and surface area (S, μm^2^) of cells were measured using Gwyddion software (version 2.49, Czech Metrology Institute, Czech Republic) designed for visualization and analysis of data obtained by scanning probe microscopy (appendix fig. 8).

### 2.5 Roughness measurement

To calculate the surface roughness, a 3×3 µm membrane surface area was scanned for each cell. Using the roughness analysis tool in Nova program the values of root-mean-square roughness (mean roughness of the surface), and maximum height difference (the difference between the highest and lowest points of the surface profile) were calculated for each frame (appendix fig. 3).

The roughness analysis tool was also used to measure the maximum cell height and calculate the average height of membrane protrusions.

### 2.6 Measurement of membrane surface structures

The number of membrane surface structures (protrusions and invaginations) was counted and their linear dimensions (height for protrusions; depth and width for invaginations) were measured to identify the topography features of the surface of the reticulocytes. To do this, the obtained scans were used to plot the profile curves of the cell surface, on which the linear dimensions were measured and the number of protrusions and invaginations was counted for each cell manually (appendix fig. 4–6).

### 2.7 Statistical analysis

The results were statistically analyzed using GraphPad Prism 8.01 software. Data distribution is presented as a scatter plot with arithmetic mean value ± SEM. The significance of the differences between the mean values due to the non–normal distribution of data according to Shapiro-Wilk test was evaluated by the Kruskal-Wallis test followed by Dunn’s post hoc test (ns – non-significant; *P<0.05; **P<0.01; ****P<0.0001).

## Results

### 3.1 AFM imaging

The images of immature reticulocytes were represented by cells of two types: with a noticeable bulge on the surface (Figure 1, A-C) and without it (Figure 1, D-F), which is supposed to be the early stage of reticulocyte maturation with the remains of the nucleus inside, and with a later one where the threads of reticulum are found throughout the entire area of the cell. The microrelief of the surface of the reticulocytes (Figure 1, G-I) was represented by protrusions (Figure 1, I1) and invaginations of irregular or rounded shape of big (Figure 1, I2) and small (Figure 1, I3) sizes.

**Figure 1.**
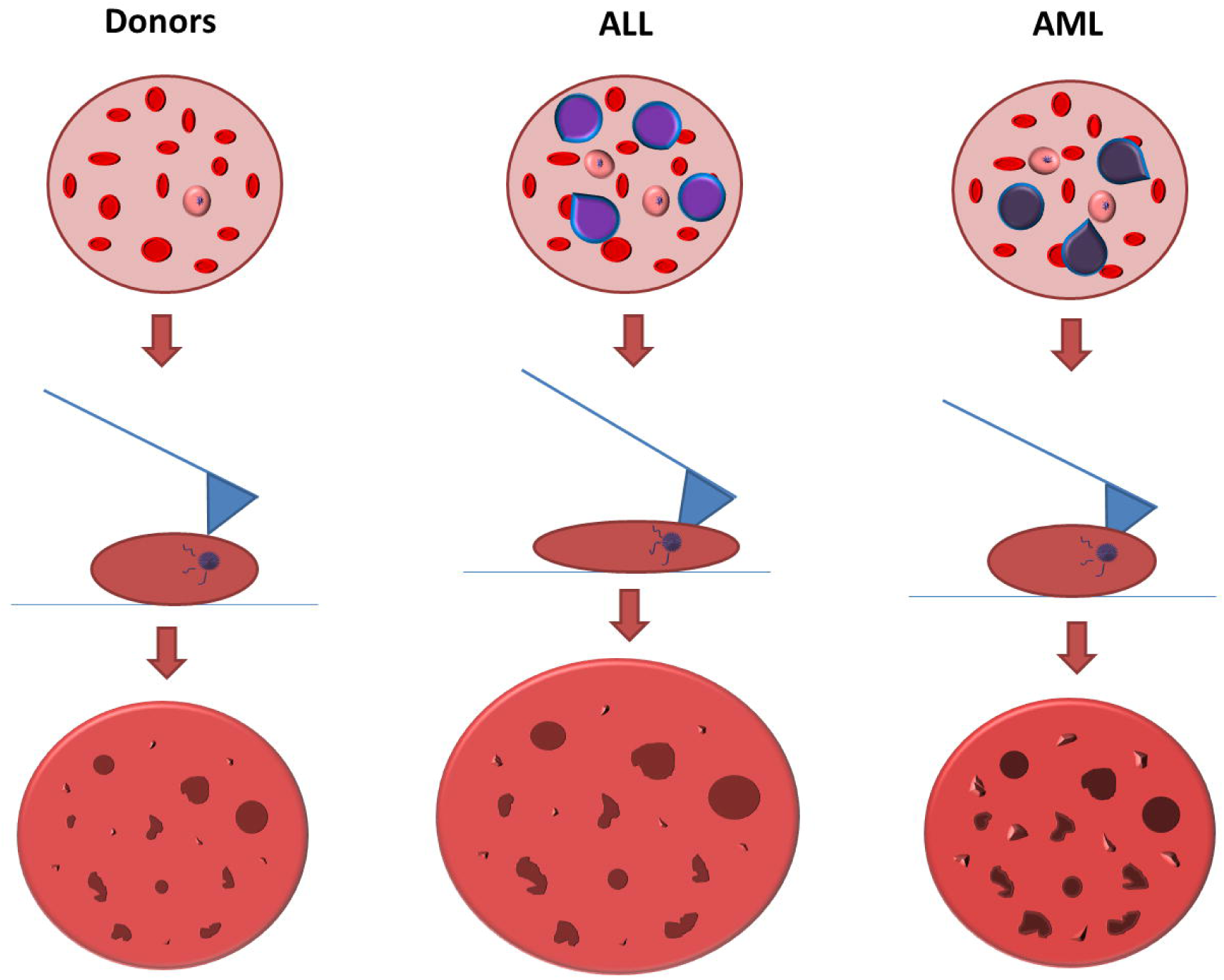
**Representative topography (contact mode: A, D, G; contact error mode: B, E, H; 3D-image: C, F, I) AFM images of immature reticulocytes: cell with a noticeable bulge on a surface (A-C), cell without a bulge (D-F), 3×3 μm cell surface site (G-I) with protrusions (I1), big (I2), and small (I3) invaginations of irregular or rounded shape.** AFM = atomic force microscopy.

A detailed study of the membrane surface of reticulocytes in areas of 3×3 μm showed that the most common invaginations of three types were: small invaginations up to 100 nm wide, rounded large invaginations of about 500 nm wide, and large shallow invaginations with an additional cavity inside (Figure 2).

**Figure 2.**
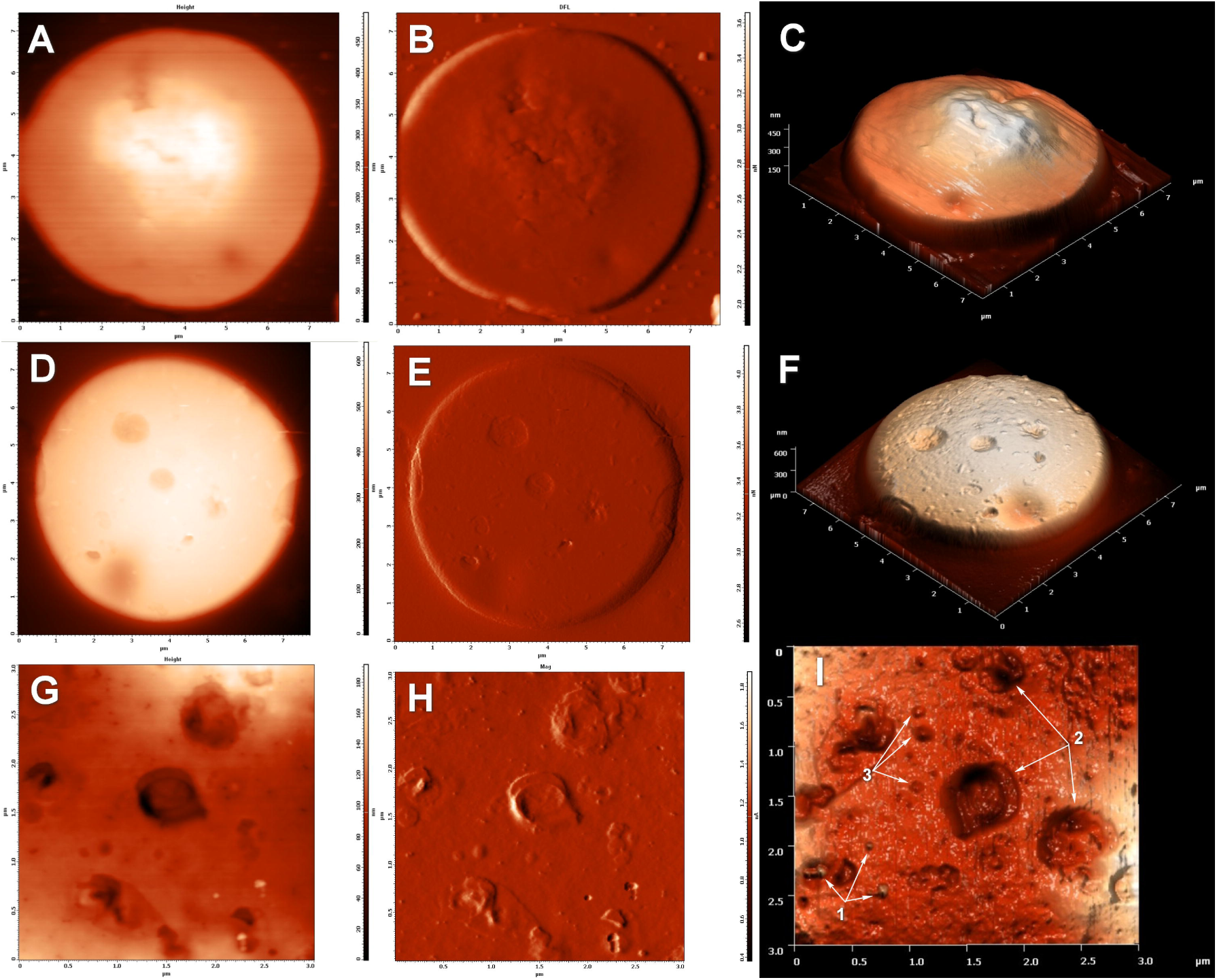
**Representative (contact mode: A, D, G; contact error mode: B, E, H; 3D-image: C, F, I) AFM images of membrane invaginations: small invaginations (A-C), big invaginations (D-F), invaginations with an additional cavity (G-I)** AFM = atomic force microscopy.

### 3.2 Young’s modulus analysis

To evaluate the stiffness of the surface of reticulocytes, Young’s modulus was measured. No significant differences between its values in groups of donors and patients with ALL and AML were found. Nevertheless, there was noted a tendency for almost one and a half decrease in membrane stiffness (Table 1, Figure 3).

**Table 1.** Average Young’s modulus of immature reticulocytes in groups of healthy donors and patients with ALL* and AML†.

| Sample | Young's modulus (mPa) | SEM (mPa) | Total no. of sampling |
| --- | --- | --- | --- |
| Donors | 5.30 | 0.76 | 72 |
| Patients with ALL | 3.94 | 0.59 | 82 |
| Patients with AML | 3.46 | 0.38 | 88 |

\* Acute lymphoblastic leukemia
† Acute myeloid leukemia

**Figure 3.**
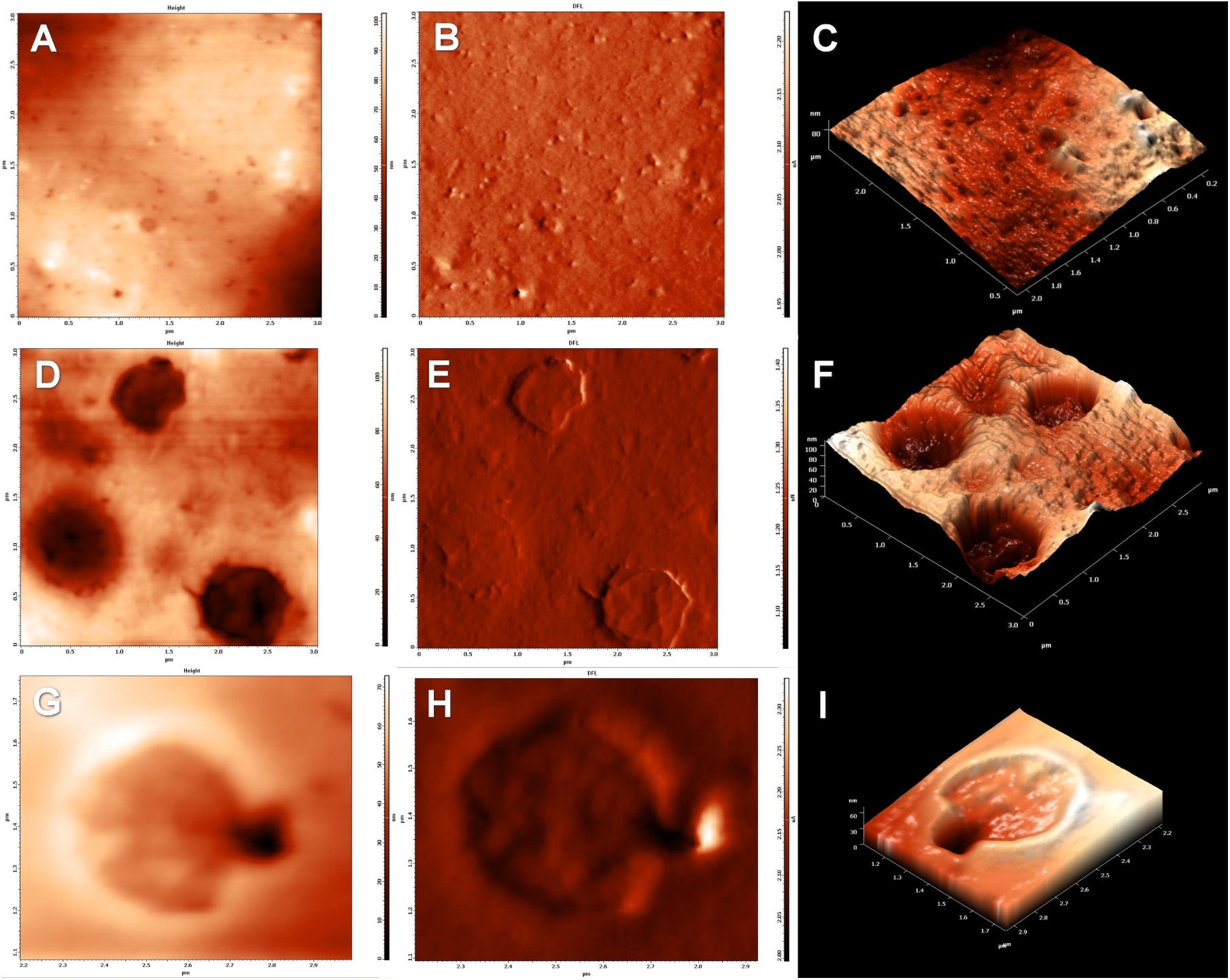
Average Young’s modulus of immature reticulocytes in groups of healthy donors and patients with ALL and AML. Dots represent individual values and whiskers represent mean value ± SEM. ALL = acute lymphoblastic leukemia, AML = acute myeloid leukemia.

### 3.3 Morphometric analysis

According to the data of morphometric analysis, it was found that in the group of patients with ALL the surface area of reticulocytes increased by 18.5% (P<0.0001), volume increased by 8.7% (P<0.05) and the height of the cells decreased by 7.8% (P<0.01) compared with the control group (Table 2, Figure 4).

**Table 2.**
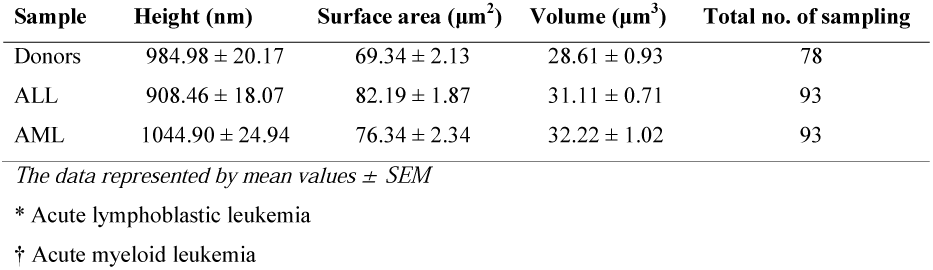
Average morphometric properties of immature reticulocytes in groups of healthy donors and patients with ALL* and AML†.

| Sample | Height (nm) | Surface area ($\mu\text{m}^2$) | Volume ($\mu\text{m}^3$) | Total no. of sampling |
| --- | --- | --- | --- | --- |
| Donors | 984.98 ± 20.17 | 69.34 ± 2.13 | 28.61 ± 0.93 | 78 |
| ALL | 908.46 ± 18.07 | 82.19 ± 1.87 | 31.11 ± 0.71 | 93 |
| AML | 1044.90 ± 24.94 | 76.34 ± 2.34 | 32.22 ± 1.02 | 93 |

*The data represented by mean values ± SEM*
\* Acute lymphoblastic leukemia
† Acute myeloid leukemia

**Figure 4.**
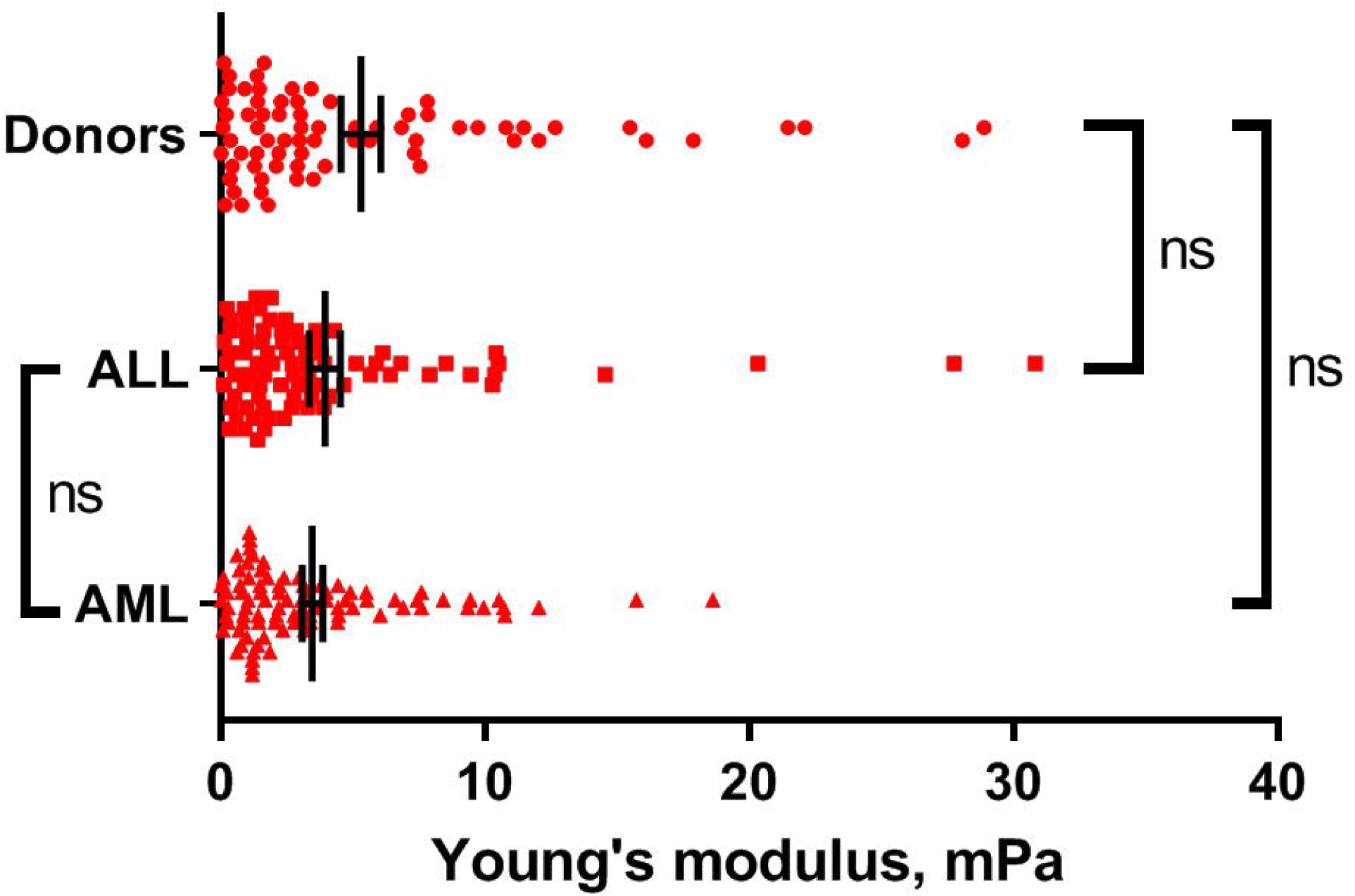
Morphometric properties of immature reticulocytes in groups of healthy donors and patients with ALL and AML. Dots represent individual values and whiskers represent mean value ± SEM. ALL = acute lymphoblastic leukemia, AML = acute myeloid leukemia.

In the group of AML patients, there was also noted a tendency to increase in surface area size and cell volume, but a significant difference was found only for volume which increased by 12.6% (P<0.01).

Also, significant differences in height (P<0.0001) and surface area (P<0.01) between groups of ALL and AML patients were found. In the group of patients with AML, the cells were 15% higher (P <0.0001) and with a 7.7% less surface area (P <0.01) than in the group with ALL.

### 3.4 Roughness analysis

For reticulocytes of patients with AML, a significant increase in the cells’ surface roughness of 35.5% (P<0.01) was established in comparison with the control group. Analyzing the maximum height difference of the surface area in the group of AML patients, there was found a significant increase of 31.9% (P<0.01) compared with the control group.

In reticulocytes from the group of ALL patients, a tendency to increase in the values of the cell surface roughness parameters was also observed, but it was not statistically significant (Table 3, Figure 5).

**Table 3.**
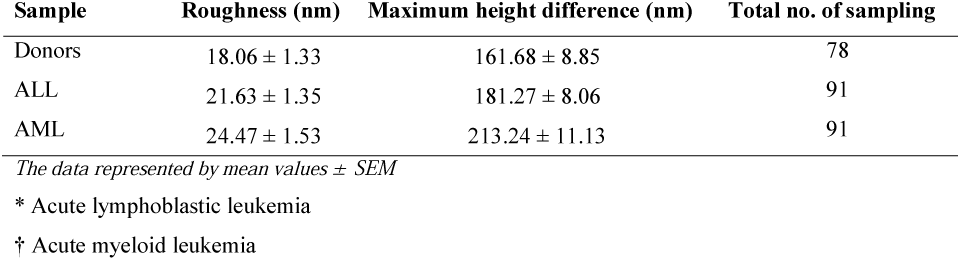
Average roughness properties of immature reticulocytes in groups of healthy donors and patients with ALL* and AML†.

**Figure 5.**
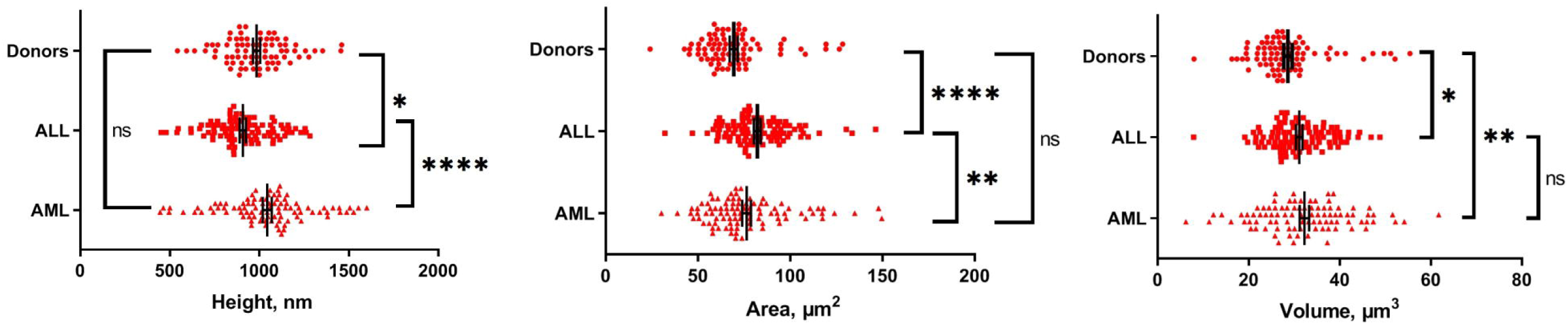
Roughness properties of immature reticulocytes surface in groups of healthy donors and patients with ALL and AML. Dots represent individual values and whiskers represent mean value ± SEM. ALL = acute lymphoblastic leukemia, AML = acute myeloid leukemia.

### 3.5 Membrane surface structures

In the group of patients with ALL, there was a significant increase in the width of invaginations of 18% (P <0.01).

In the group of patients with AML, there was a significant increase in the height of protrusions of 36.2% (P <0.01), the depth of invaginations of 24.8% (P <0.01), and their width of 18.2% (P <0.01).

There was also a difference between patients with ALL and AML revealing a significant (P <0.01) difference between the number of protrusions on the cells. In the AML group, there were 42.8% more of them than in the ALL group (Tables 4, 5; Figures 6, 7).

**Table 4.** Average properties of protrusions in membranes of immature reticulocytes in groups of healthy donors and patients with ALL* and AML†.

| Sample | Number | Total no. of sampling | Height (nm) | Total no. of sampling |
| --- | --- | --- | --- | --- |
| Donors | 25.51 ± 2.20 | 77 | 99.70 ± 5.84 | 79 |
| ALL | 17.79 ± 1.15 | 82 | 107.09 ± 6.15 | 91 |
| AML | 25.41 ± 1.68 | 69 | 135.80 ± 8.92 | 91 |

*The data represented by mean values ± SEM*
\* Acute lymphoblastic leukemia
† Acute myeloid leukemia

**Table 5.** Average properties of invaginations in membranes of immature reticulocytes in groups of healthy donors and patients with ALL* and AML†.

| Sample | Number | Total no. of sampling | Width (nm) | Depth (nm) | Total no. of sampling |
| --- | --- | --- | --- | --- | --- |
| Donors | 13.18 ± 1.10 | 76 | 475.33 ± 23.61 | 27.10 ± 2.19 | 76 |
| ALL | 10.56 ± 0.56 | 91 | 561.13 ± 24.70 | 27.64 ± 1.29 | 91 |
| AML | 10.16 ± 0.72 | 90 | 561.73 ± 20.36 | 33.81 ± 1.84 | 89 |

*The data represented by mean values ± SEM*
\* Acute lymphoblastic leukemia
† Acute myeloid leukemia

**Figure 6.**
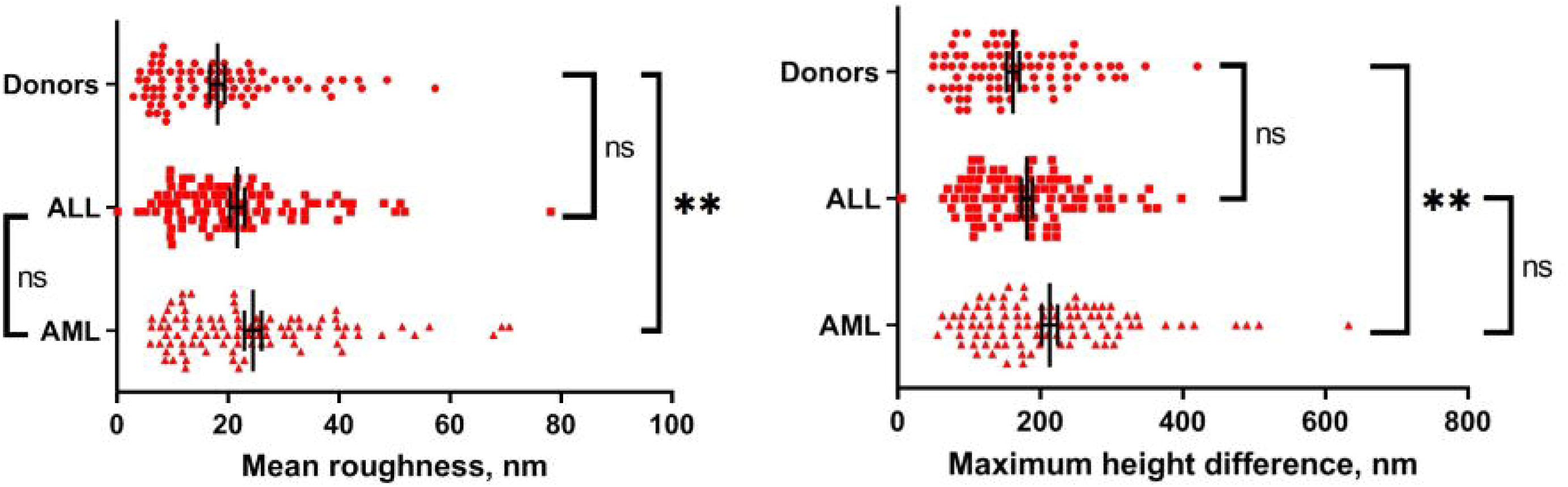
Properties of protrusions in groups of healthy donors and patients with ALL and AML. Dots represent individual values and whiskers represent mean value ± SEM. ALL = acute lymphoblastic leukemia, AML = acute myeloid leukemia.

**Figure 7.**
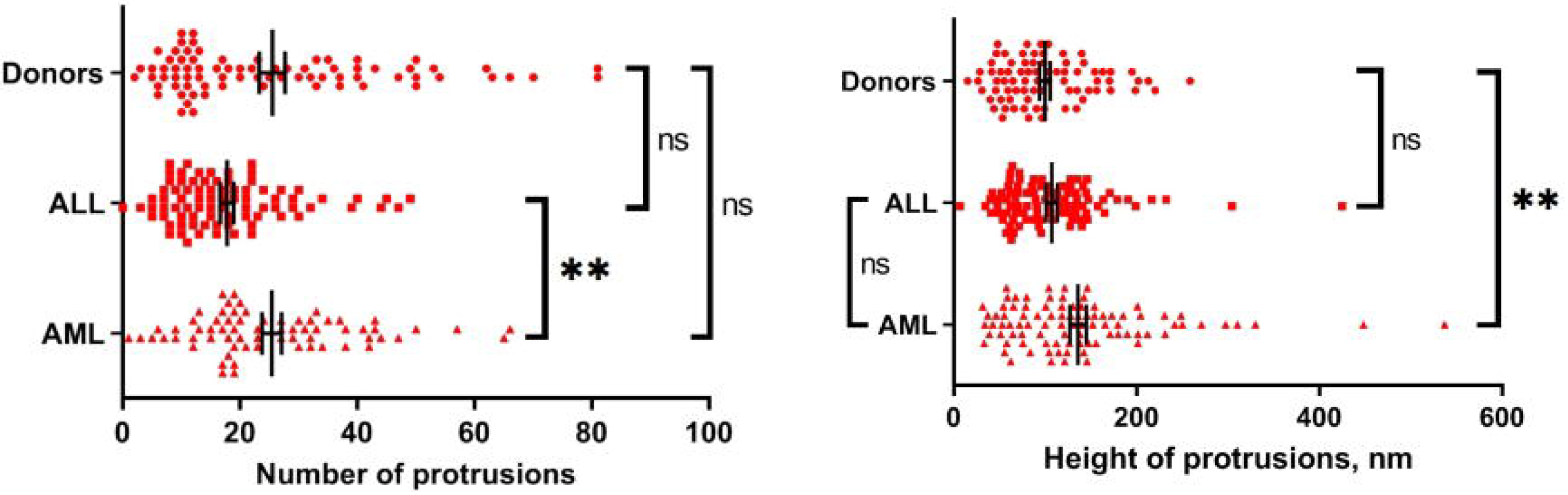
Properties of invaginations in groups of healthy donors and patients with ALL and AML. Dots represent individual values and whiskers represent mean value ± SEM. ALL = acute lymphoblastic leukemia, AML = acute myeloid leukemia.

## Discussion

This work shows that the bloodstream of both ALL and AML patients contain immature reticulocytes with an increased cell volume and, in ALL patients exclusively, reticulocytes with an increased surface area and decreased cell height.

As shown in the research, the plasma membrane surface of immature reticulocytes in AML patients is characterized by noticeable roughness. The width and depth of invaginations and the height of protrusions are increased compared to the control group. The surface relief of immature reticulocytes in patients with acute lymphoblastic leukemia is characterized by invaginations with increased width. I also noted a tendency towards a decrease in membrane stiffness in both groups of patients with both ALL and AML, requiring further studies with an increase in the number of samples to confirm or refute the observed results.

Comparison of the ALL and AML groups with each other showed that the cells have a larger area and smaller height in the ALL group. In the contrast, they have a smaller area and a larger height in the AML group, and a greater number of protrusions on the membrane, which could be associated with different mechanisms of compensatory-adaptive reactions in ALL and AML and also requires further studies.

Comparing data obtained in this research with data from the work of Gifford et al., it can be noted that the surface area and volume of reticulocytes presented in their work are greater that in the presented research^18^.

I could attribute this discrepancy to different methodological approaches since the authors of the referenced research used a microchannel device to measure the cell size, which makes it possible not to perform special sample preparation before use. In the presented study, the blood smears were air-dried immediately before scanning with an atomic force microscope, which led to a loss of cell mass due to water evaporation, and a decrease in their size, while maintaining their shape.

Deformability is important in the regulation of the release of reticulocytes from bone marrow to the blood^19^. The tendency of the reticulocyte membrane stiffness to decrease is observed in both groups of patients with acute lymphoblastic and acute myeloid leukemia probably act as a compensatory-adaptive mechanism that allows cells to migrate through the vascular endothelium more easily. There is evidence of an increased incidence of cardiovascular group diseases in individuals who survived acute lymphoblastic leukemia^20-21^. As additional factors causing increased stiffness and endothelial dysfunction, researchers indicate some chemotherapy regimens involving the use of anthracycline drugs (doxorubicin, daunomycin and others)^22-23^. It has been proven that the action of drugs that cause endothelium-dependent relaxation has a cytotoxic effect on vascular endothelial cells, especially in small-diameter vessels^24^.

Leukemia cells (blasts) inhibit proliferation of normal cell lines and causes immature cells with bigger sizes to enter the bloodstream, while normally they mature in the bone marrow^25^. Myelodysplastic syndromes and some types of anemia cause an increase in reticulocytes volume^26^. This is consistent with the presented data on the increased size of immature reticulocytes in the groups of patients with ALL and AML.

An increase in the width and depth of invaginations in the cell membrane and increasing roughness values possibly indicate an intensification of the processes of endo- and exocytosis accompanying reticulocytes maturing into erythrocytes — a process when immature cells lose part of their membrane and the remains of intracellular organelles^18,27^. It has been shown that immature reticulocytes have an increased ability to extrude oxidized waste through exo- and endovesiculation processes compared to mature ones^8^. Observed membrane invaginations could point to a model of exocytosis of autophagocytosed cytoplasmic content during reticulocyte maturation, according to which autophagosomes combine with endosomes to form large, autophagic compartments that fuse with the plasma membrane to release their contents by exocytosis^28-29^. In this case, small invaginations could be correlated with cadherin-based endocytosis and larger ones — with different stages of exocytosis or microvesiculation processes development.

A limitation of this study is that there were small groups of 10 persons (10 donors, 10 patients with ALL and 10 patients with AML) to perform the research. Additionally, to distinguish immature reticulocytes I used dyeing with the following observation with optical microscopy which is less accurate in comparison to automatic methods such as flow cytometry. Another important limitation of this study is that identification of protrusions and invaginations of each cell surface and measuring their linear dimensions were performed manually, however, automatic methods of this type of work do not exist.

These results demonstrate the usefulness of various atomic force microscopy techniques for detecting non-obvious morphofunctional changes in immature reticulocytes affected by tumor blasts in the bone marrow. AFM imaging makes it possible to quantitatively evaluate the features of the plasma membrane surface microrelief and the linear dimensions of cells and to measure its stiffness. These are promising tools for better understanding the dynamics of initiation and course of malignant processes in the blood system, as well as for assessing the effectiveness of their treatment. Further studies should develop the cell surface studying with a more accurate division of the reticulocyte fraction which could help distinguish specific membrane changes occurring in different conditions.

## Supporting information

Supplementary images and descriptions of methods

## Appendix. Supplementary data

The supplementary data related to this article include 17 figures with additional AFM images of cell surface and a step-by-step description of applied methods.

## Acknowledgment

The author would like to thank Mrs. Marina Yu. Skorkina from the Medicine Institute of the Belgorod State University for the research supervision, Mr. Alexey B. Fedosin for the grammar and spell check of the text, Miss Elena Shamray from the Faculty of Biology and Chemistry of the Belgorod State University for the valuable advices and Mrs. Olga V. Cherkashina from the Belgorod regional clinical hospital for the curation of the blood donation.

## Abbreviations

AFM: atomic force microscopy
ALL: acute lymphoblastic leukemia
AML: acute myeloid leukemia
IRF: immature reticulocyte fraction
EDTA: ethylenediaminetetraacetic acid
SEM: standard error of the mean

---

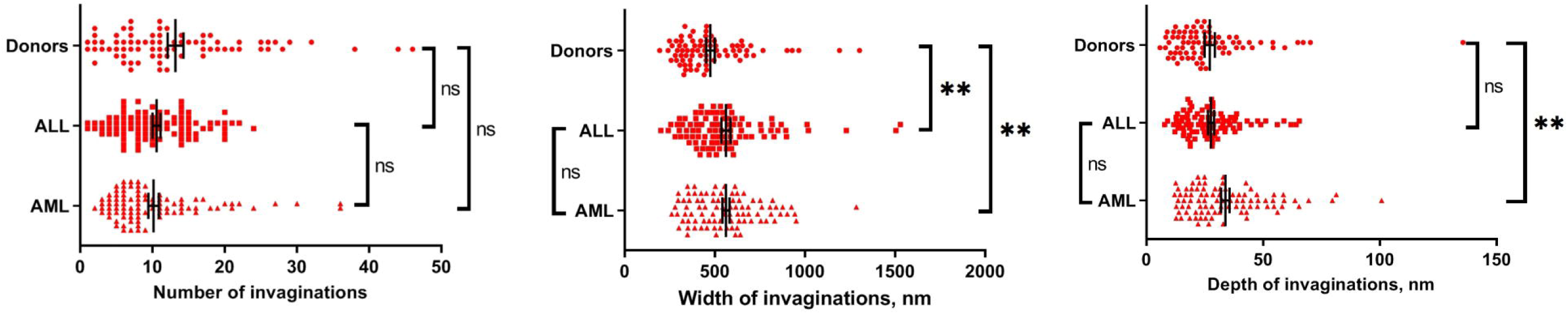

## Notes

http://dx.doi.org/10.17632/54myng9d2t.2

### Competing Interest Statement

The authors have declared no competing interest.

