## Supplementary images and descriptions of methods for "Morphometric and elastic properties of immature reticulocytes in health and during acute lymphoblastic and acute myeloid leukemia"

**Supplementary data**

**Materials and methods**

**Atomic force microscopy**

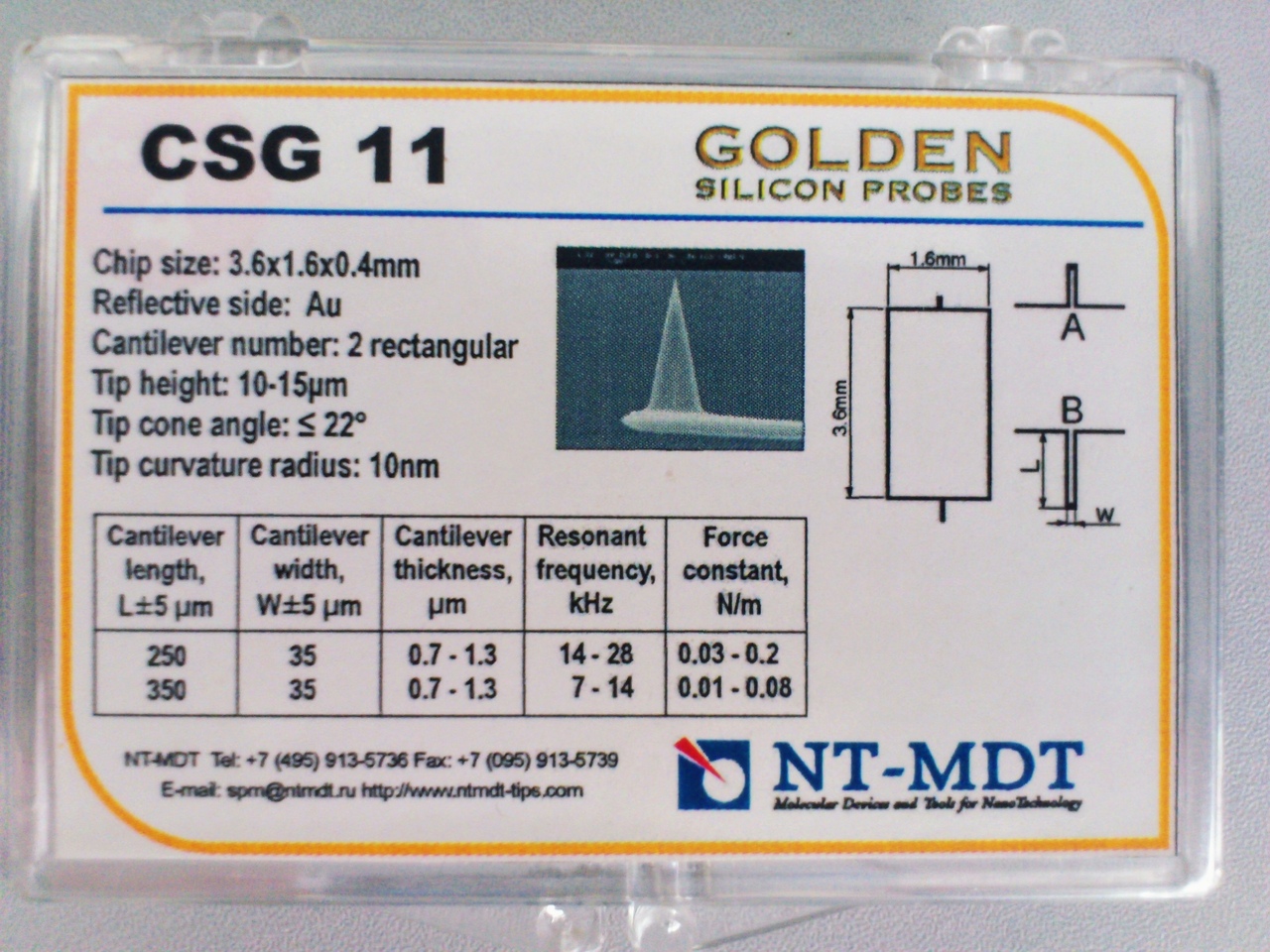

Figure 1. The set of AFM probes we used in work

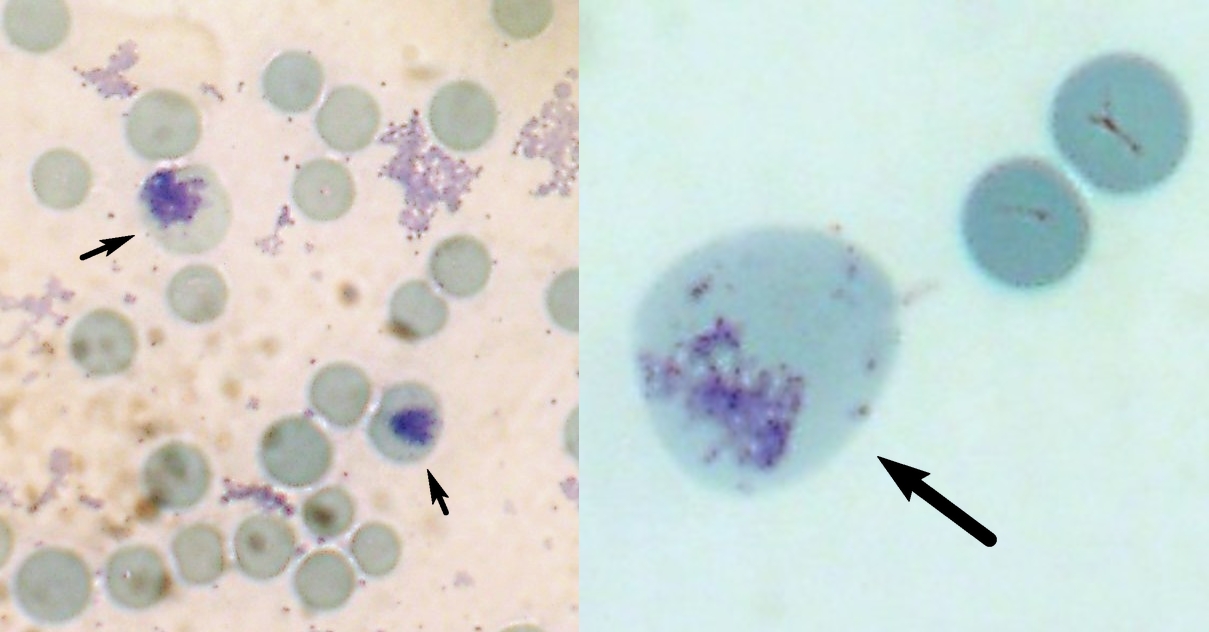

Figure 2. Examples of immature reticulocytes we chose for scanning (magnification 100x)

**Roughness analysis**

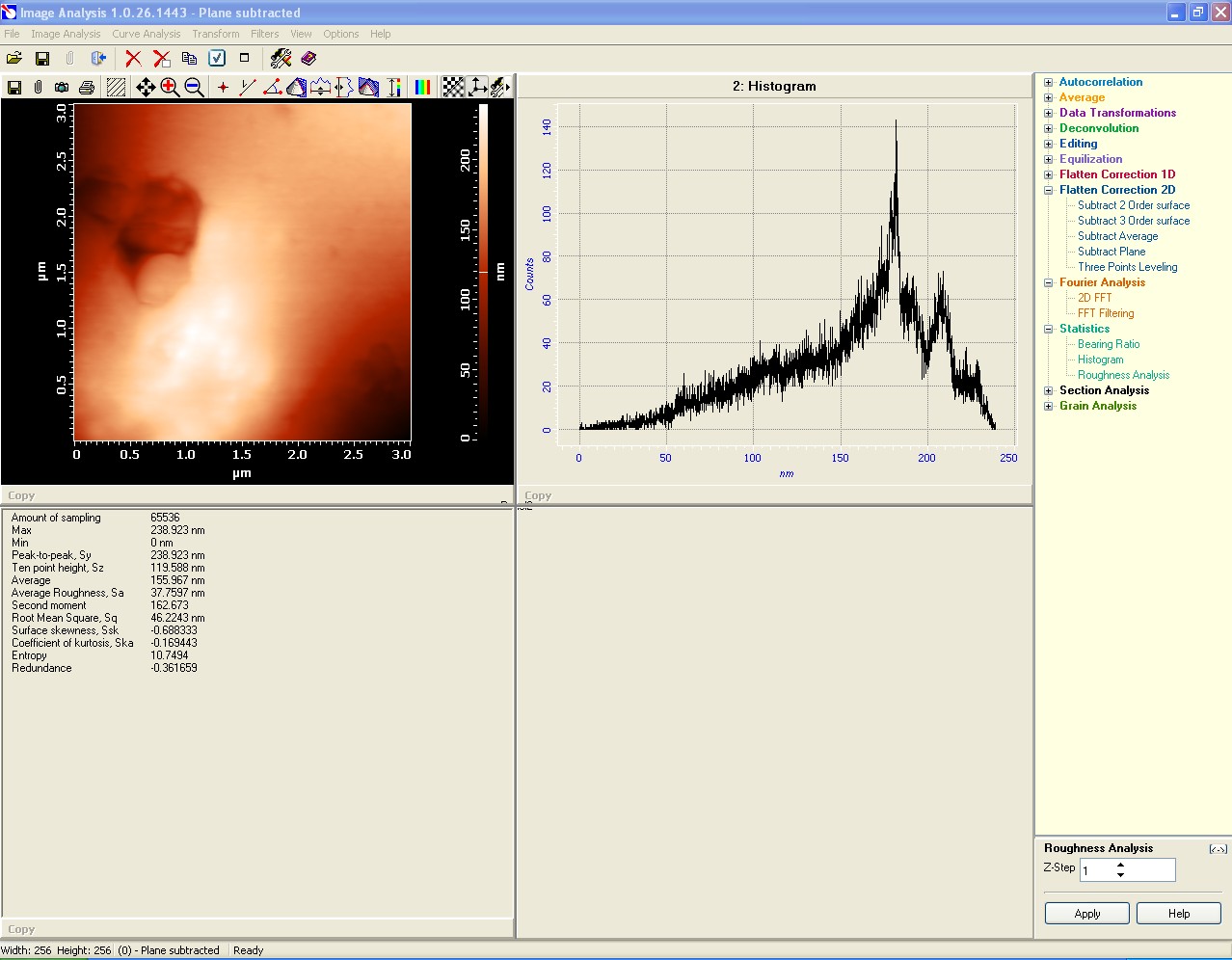

Figure 3. Application window of Roughness analysis tool from Nova software

Steps:

1. Launch Nova.

2. Open the **.mdt** file with your AFM data.

3. Select the frame for analysis.

4. Click the “Analysis” button at the top of the chosen frame.

5. Then go to “Flatten correction 2D” → “Subtract plane” → “Apply” from the right menu if you didn’t do this in the scanning process before.

6. Switch to an image with a subtracted plane by clicking on it.

7. Go to “Statistics” → “Roughness analysis” → “Apply” from the right menu.

8. Check the roughness values and cell height (“Max”) from the application window below.

Mean roughness stands for arithmetic mean roughness characterizing the arithmetic mean deviation of the values of peaks and troughs on the profile from the midline. The lower the Sa value, the smoother the surface.

Maximum height difference stands for the maximum difference in heights between the highest and lowest points of the profile surface. This parameter corresponds to the thickness of the surface layer enclosed between the planes passing through the lowest and the highest points of the surface. The solid layer lies below this layer. Thus, maximum height difference can be considered as a parameter that characterizes the thickness of the disturbed layer in which the relief changes.

**Membrane structures analysis**

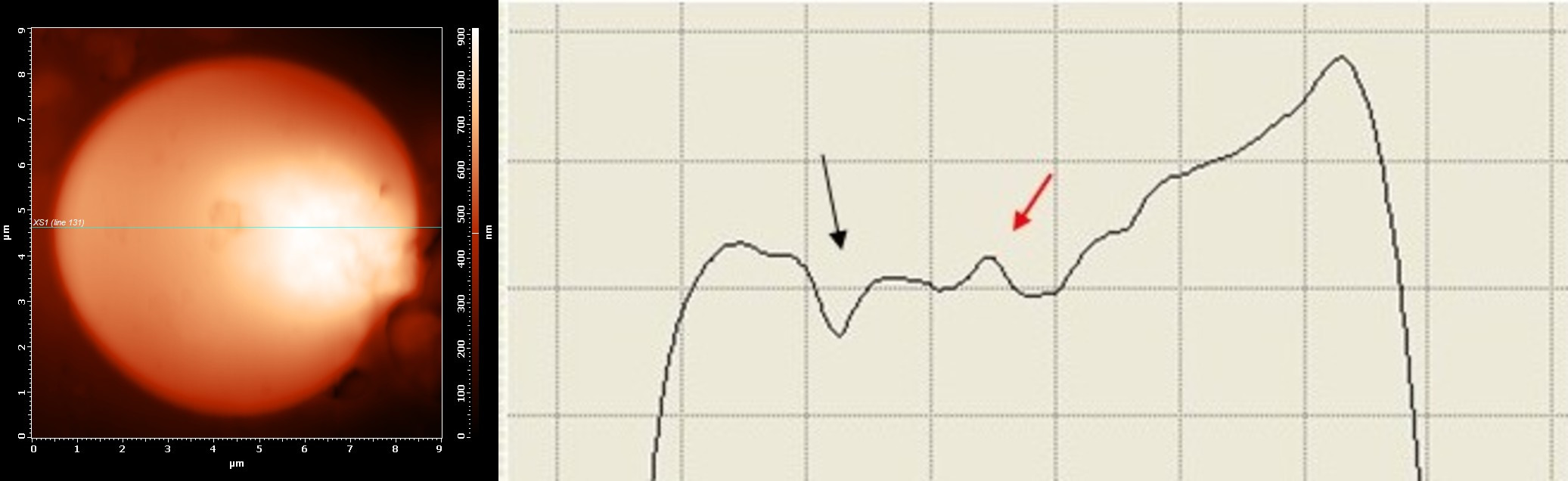

Figure 4. Protrusion (red arrow) and invagination (black arrow) on the cell membrane. The left image is a 2D scan of a cell surface, the right image – a profile curve

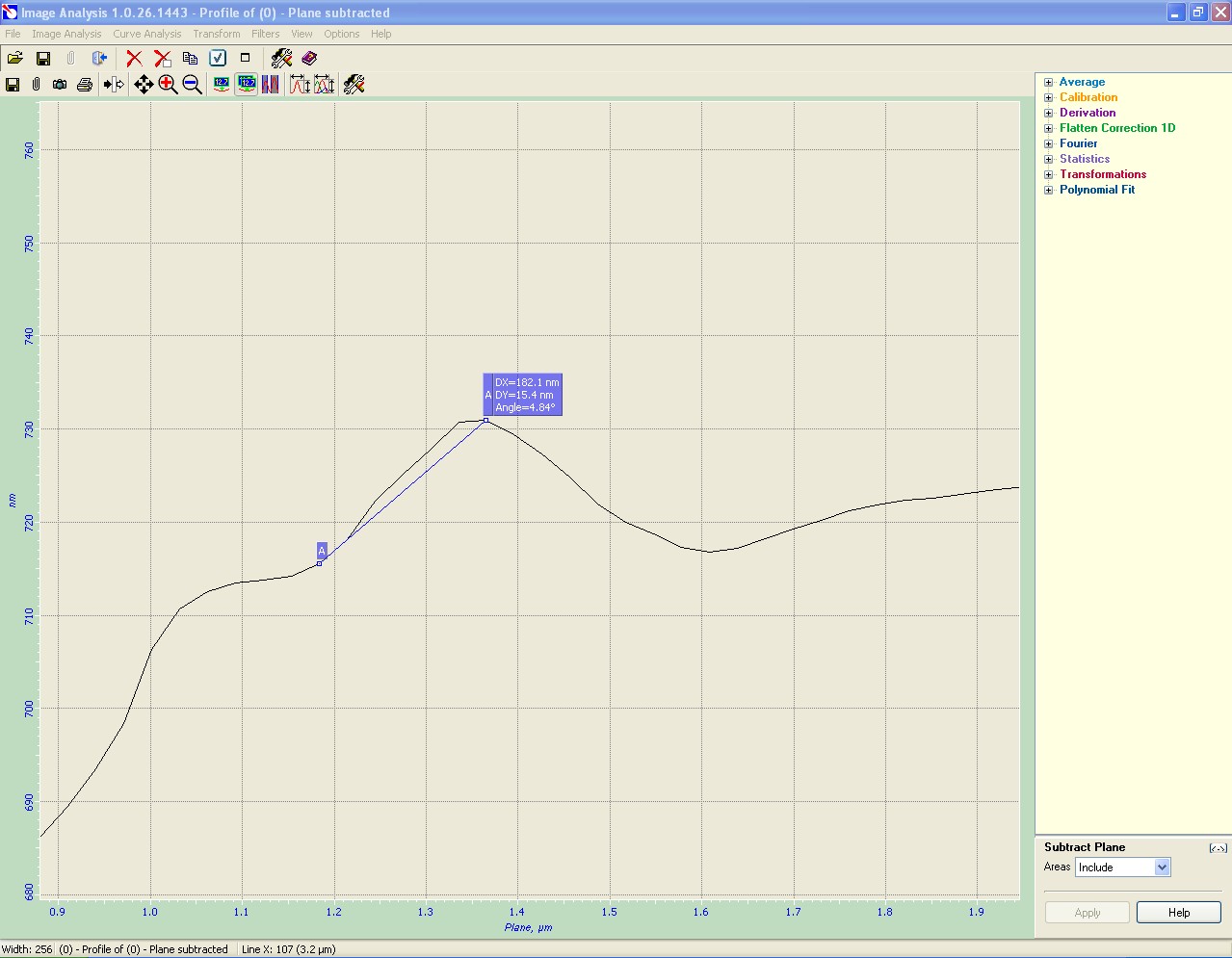

Figure 5. An example of measuring the protrusion height

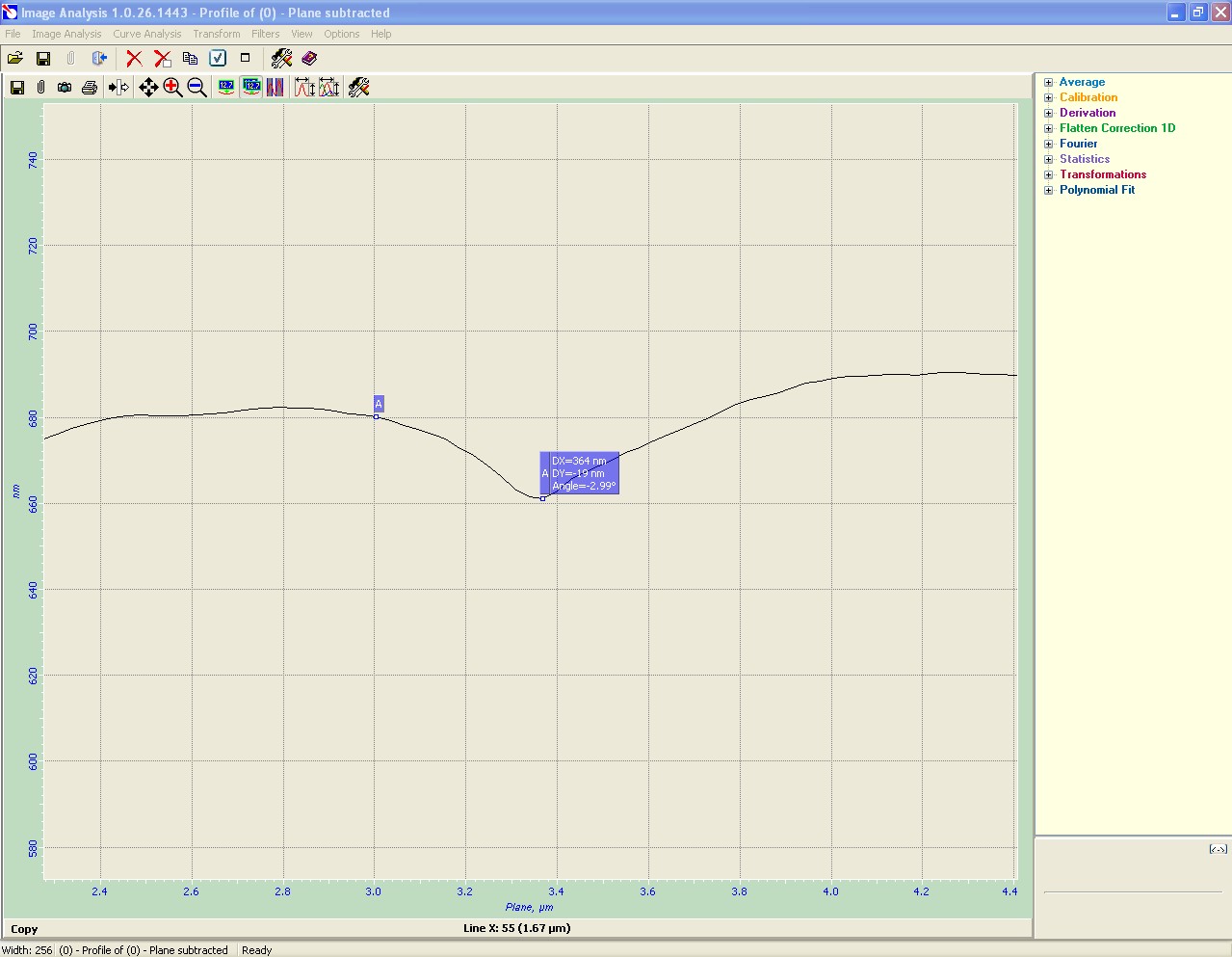

Figure 6. An example of measuring the invagination depth

Steps:

1. Launch Nova.

2. Open the .mdt file with your AFM data.

3. Select the frame for analysis.

4. Click the “Analysis” button at the top of the chosen frame.

5. Then go to “Flatten correction 2D” → “Subtract plane” → “Apply” from the right menu if you didn’t do this in the scanning process.

6. Select an image with a subtracted plane by clicking on it.

7. Go to “Section analysis” → “X section”.

8. Find protrusions or invaginations by moving the lines of the image profile.

9. Select the right window with the profile curve.

10. Select “Pair markers” above the curve.

11. Place the markers to measure the linear dimensions of membrane structure (A on the left side and B on the right one).

12. Check the “DX” value.

**Young’s modulus analysis**

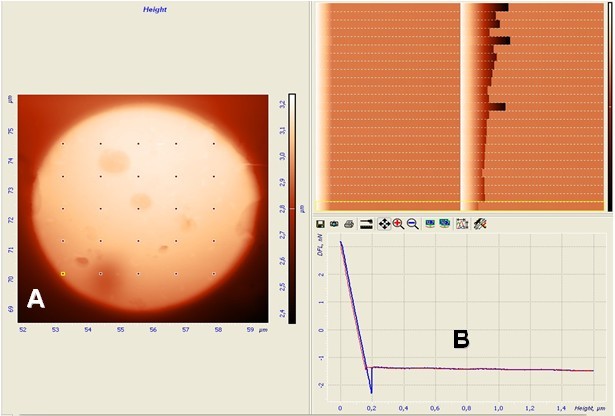

Figure 7. A – 25 local points of force apposition on cell surface, B – force curves for each point

**Area and volume measurement**

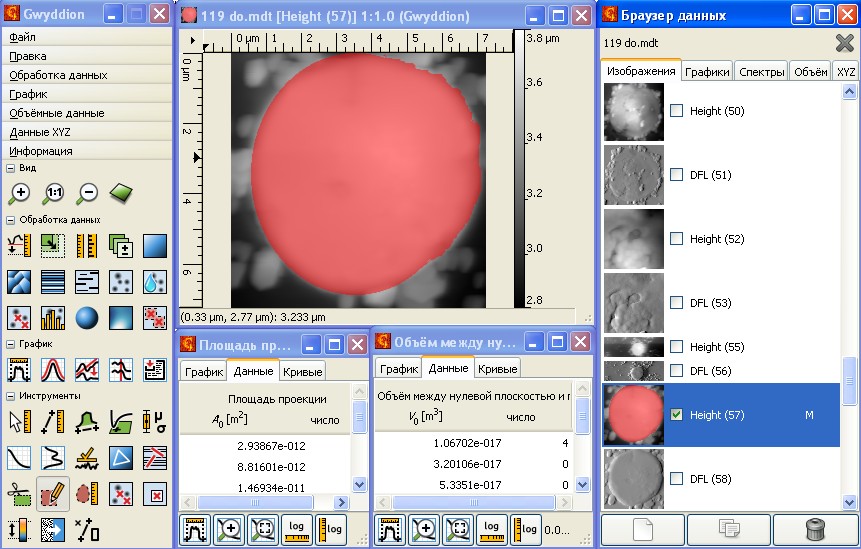

Figure 8. Application windows of Gwyddion software

Steps:

1. Launch Gwyddion.

2. Open the .mdt file with your AFM data.

3. Go to “Information” → “Data browser”.

4. Select the frame for analysis.

5. Go to “Data Process” → “Level” → “Fix Zero”.

6. Go to “Data Process” → “Level” → “Plane Level”.

7. Go to “Data Process” → “Level” → “Facet Level”.

8. Go to “Data Process” → “Level” → “Align lines”.

9. Go to “Data Process” → “Level” → “Remove horizontal scratches”.

10. Go to “Instruments → “Mask editor tool→ “Pencil”.

11. Then circle the cell precisely.

12. Go to “Instruments → “Mask editor tool→ “Fill”.

13. Fill the circled area.

14. Go to “Data processing → “Grain analysis”.

15. Select “Area” (projection area and surface area).

16. Select “Volume”.

17. Check the values (Area of the cell surface = Projection area + surface area).

**Results**

**AFM imaging**

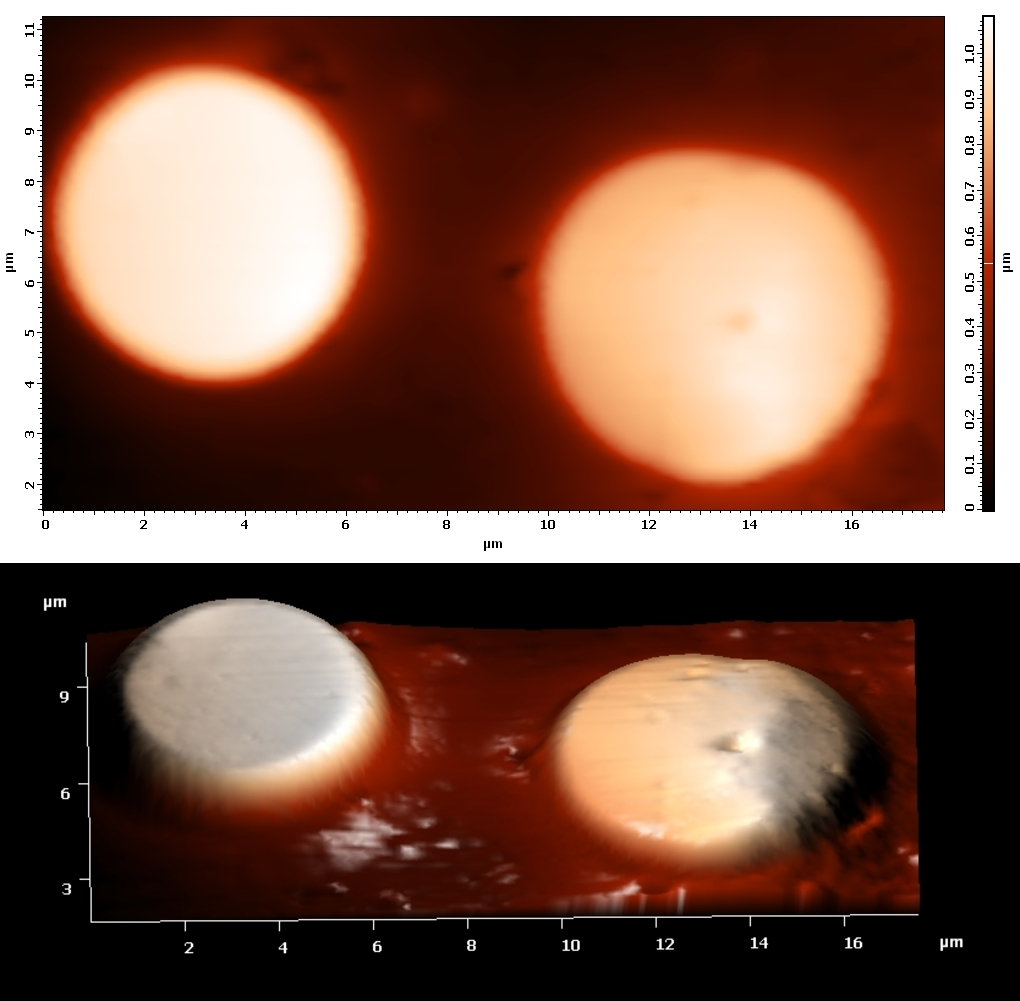

Figure 9. Immature reticulocyte (right cell) and erythrocyte (left cell) from the donor sample on the AFM images (contact mode)

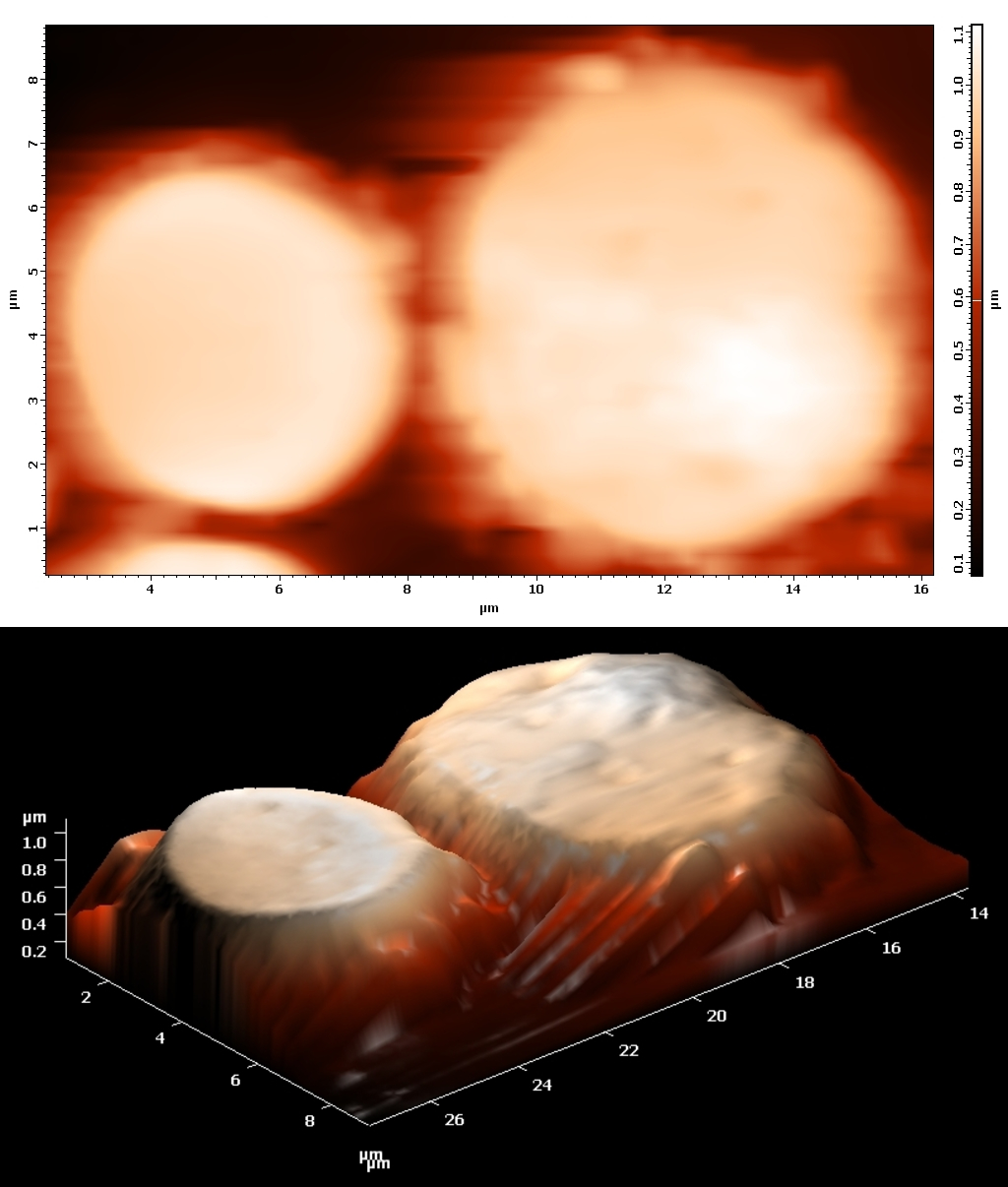

Figure 10. Immature reticulocyte (right cell) and erythrocyte (left cell) from the ALL sample on the AFM images (contact mode)

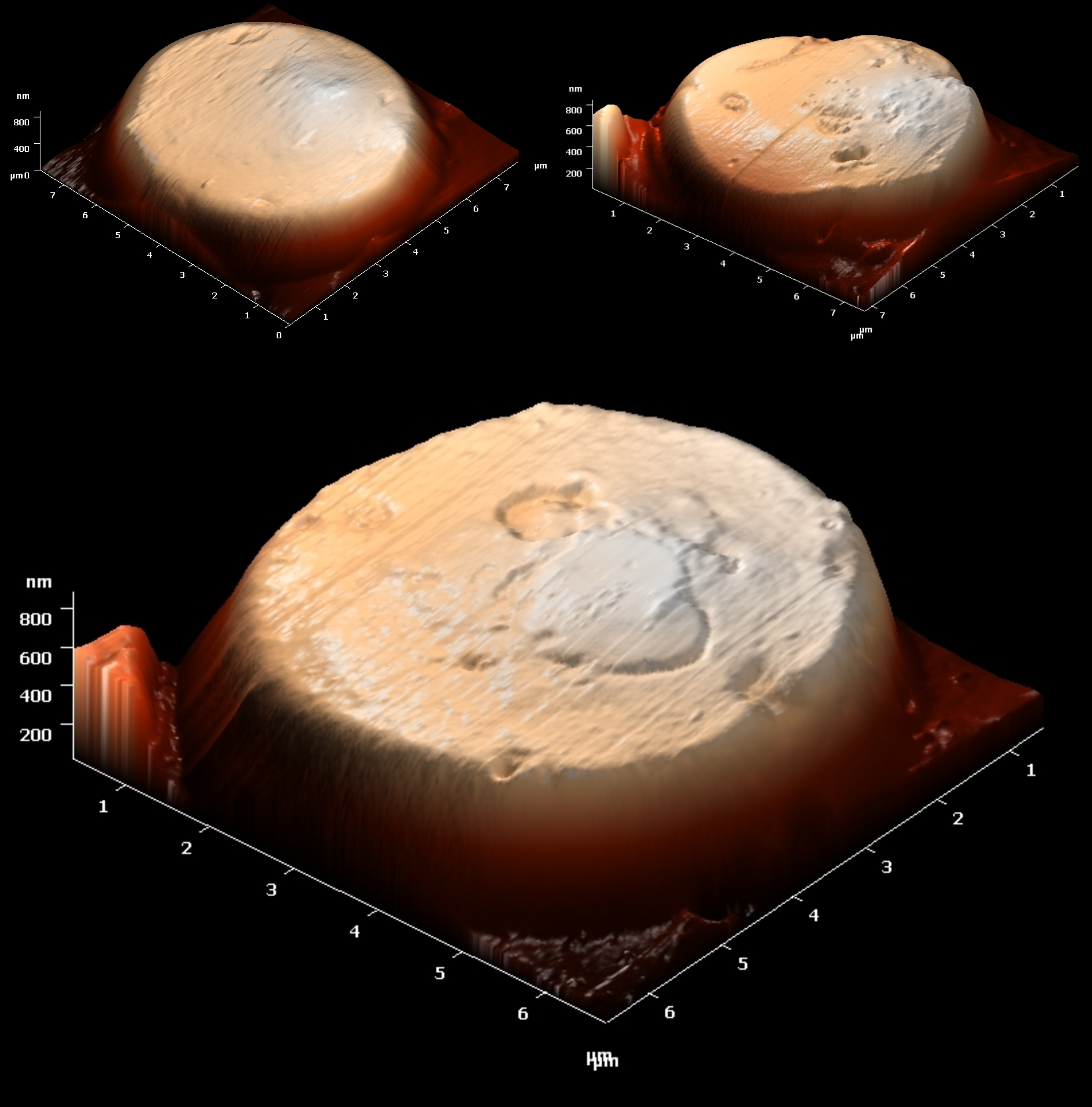

Figure 11. Examples of immature reticulocytes with bulges from donor samples

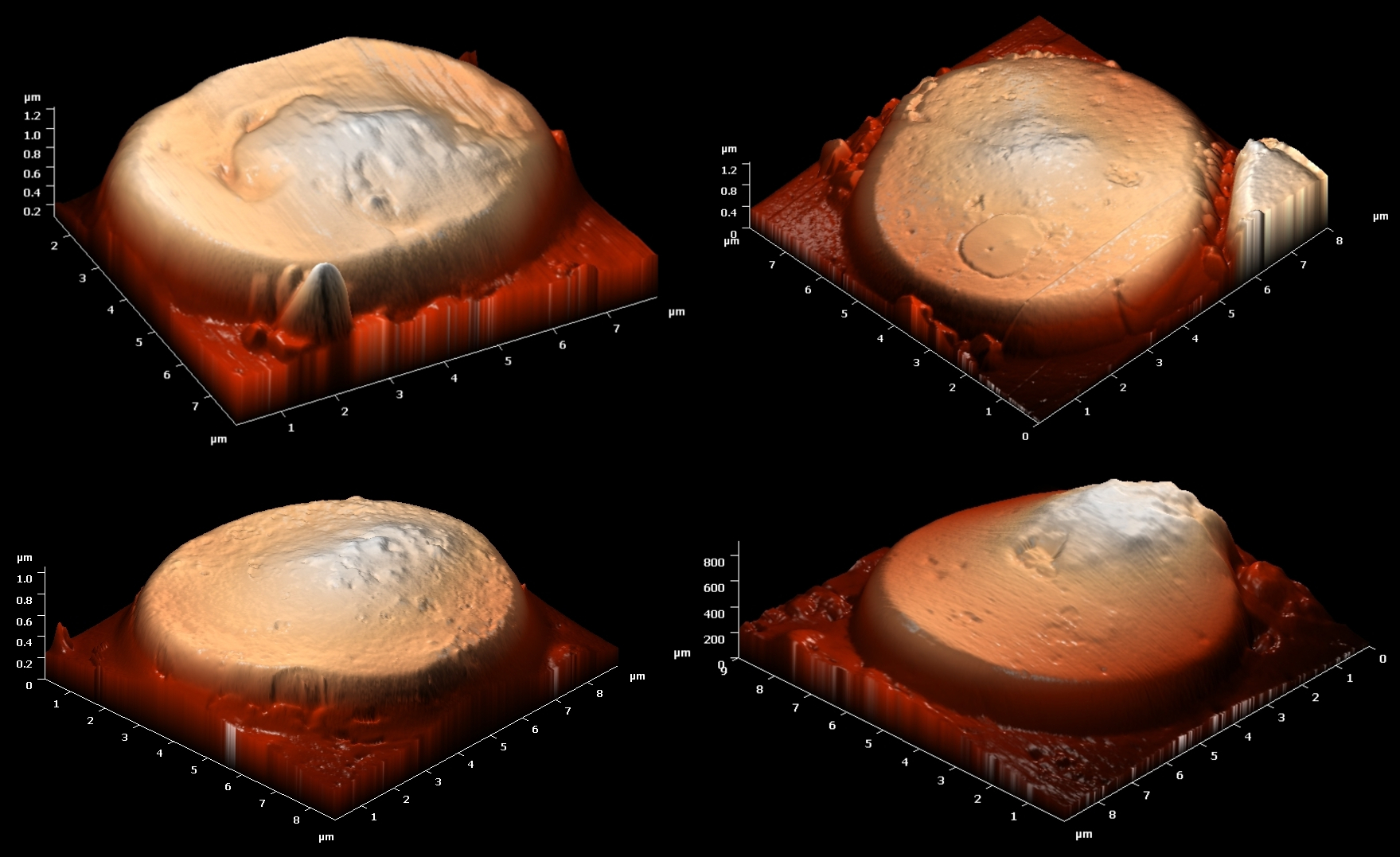

Figure 12. Examples of immature reticulocytes with bulges from ALL samples

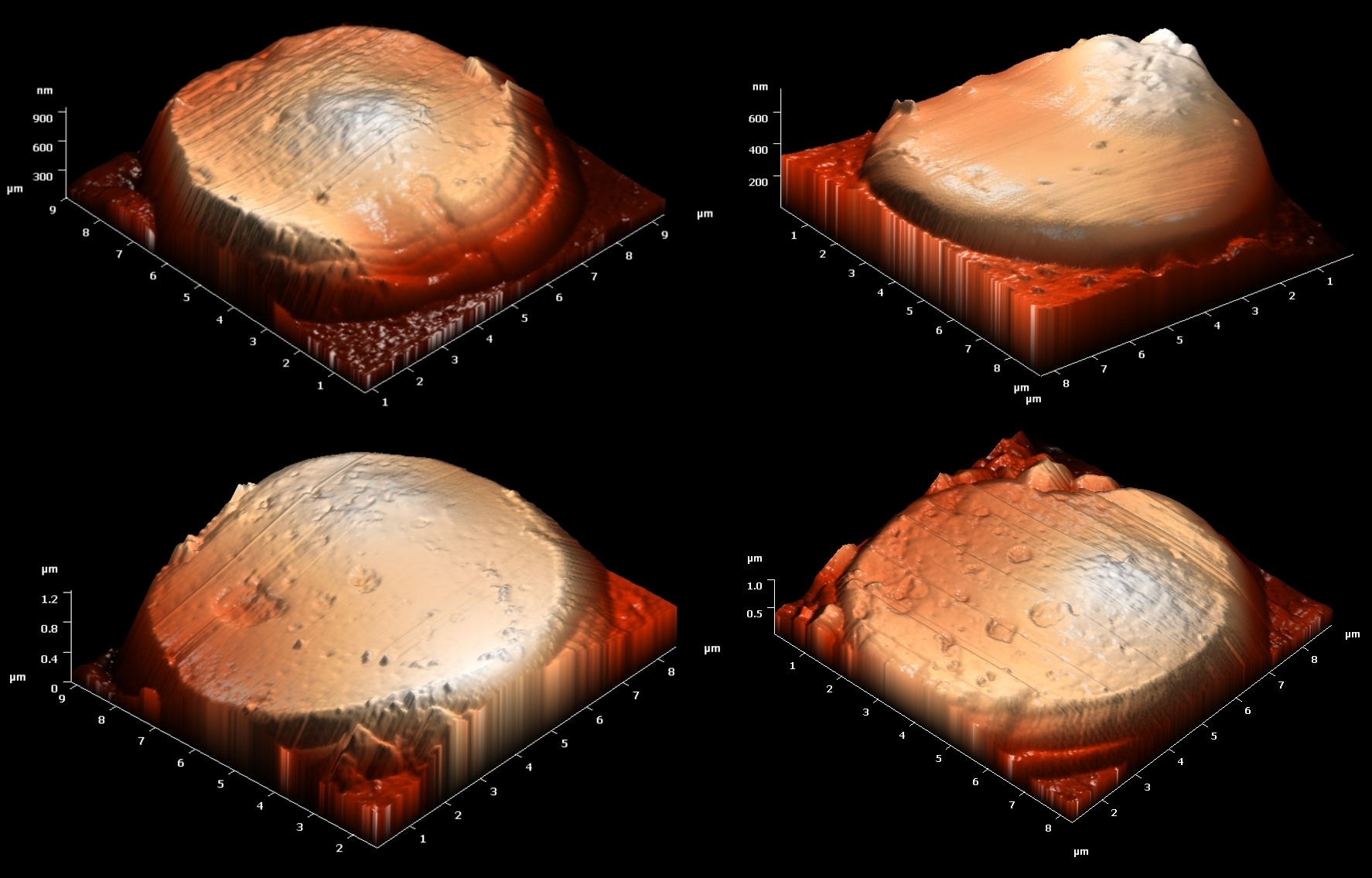

Figure 13. Examples of immature reticulocytes with bulges from AML samples

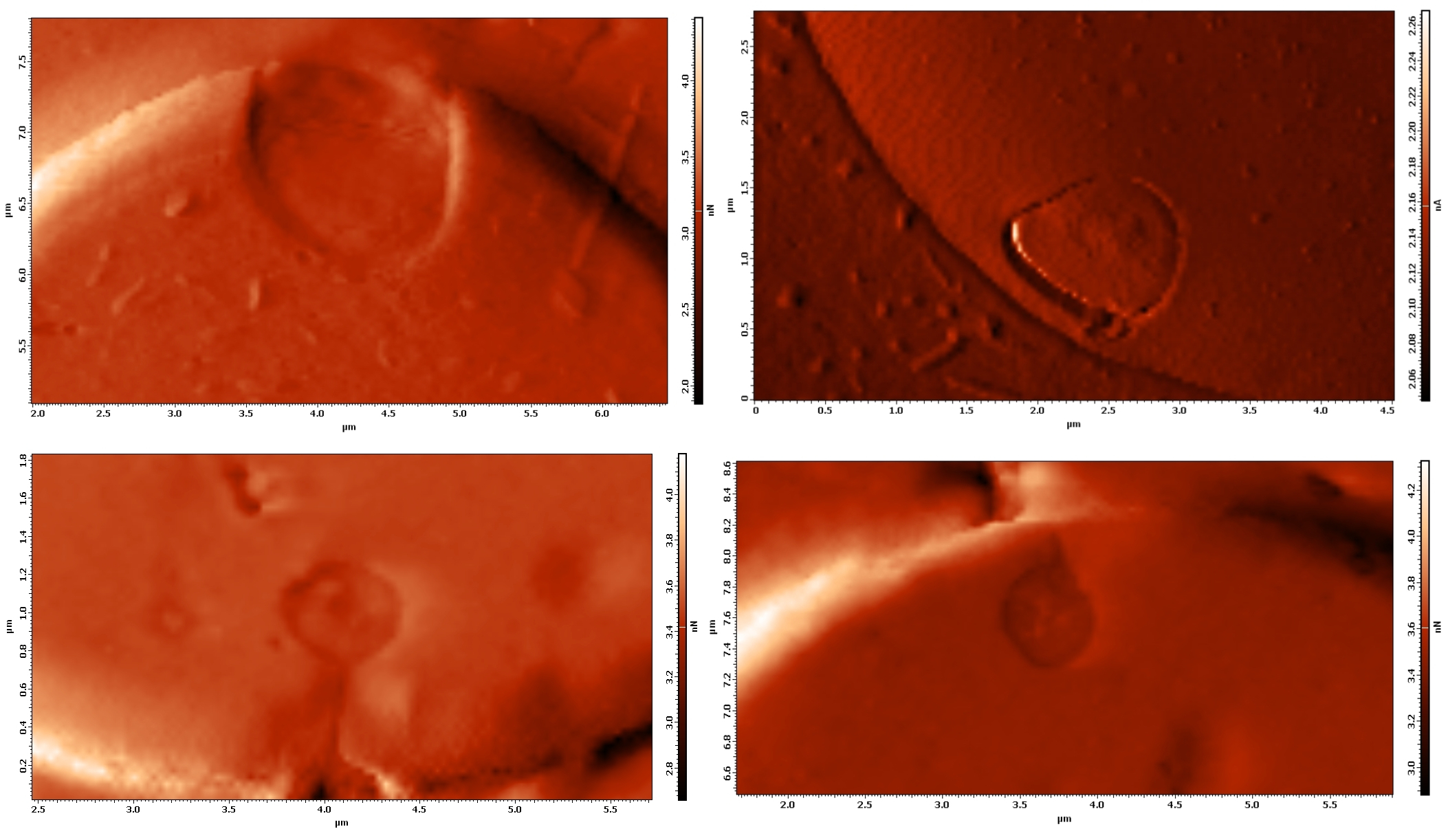

Figure 14. Examples of supposed exocytosis process from donor samples (scanned in contact error mode)

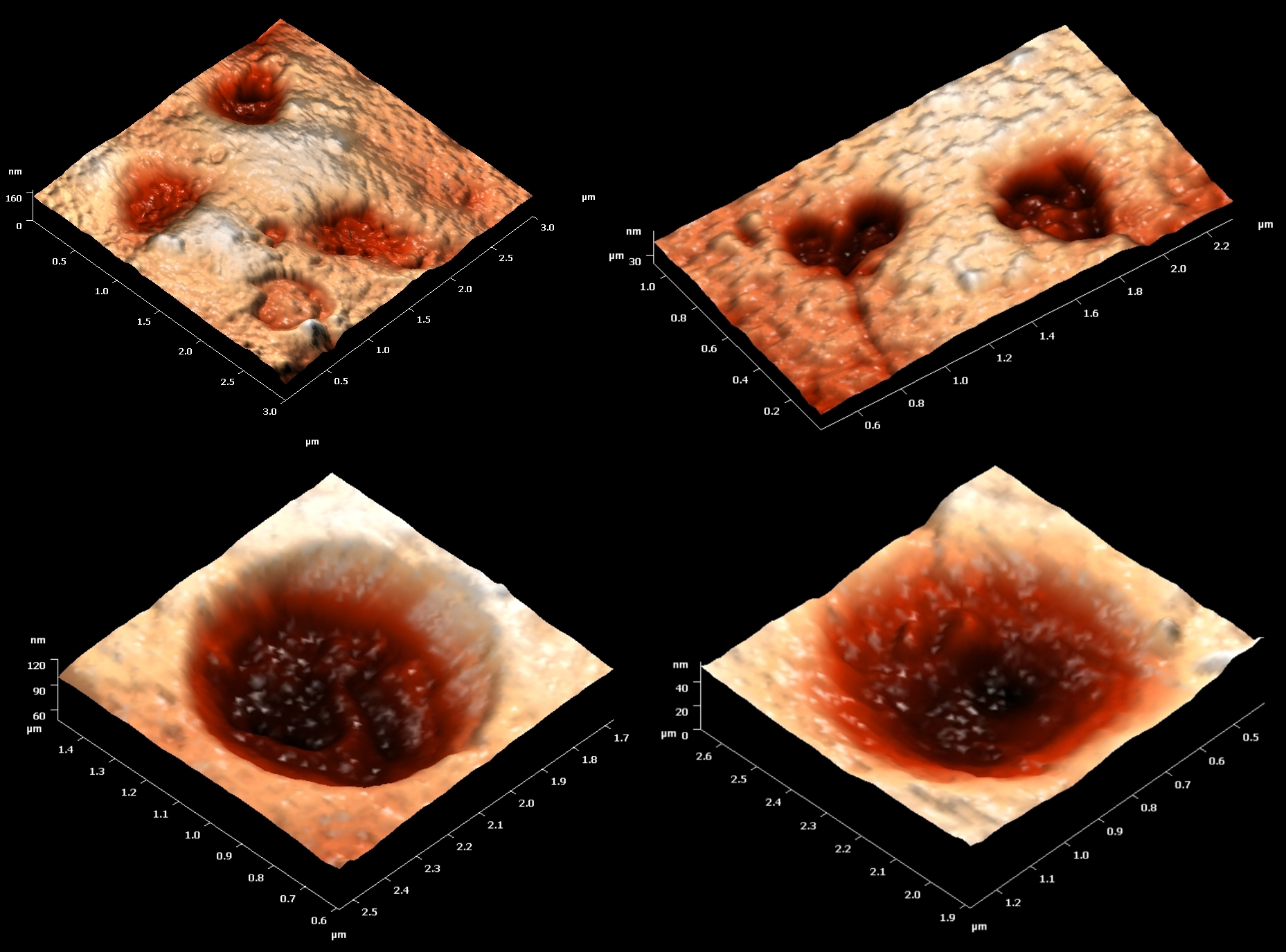

Figure 15. Examples of big invaginations from donor samples (scanned in contact mode)

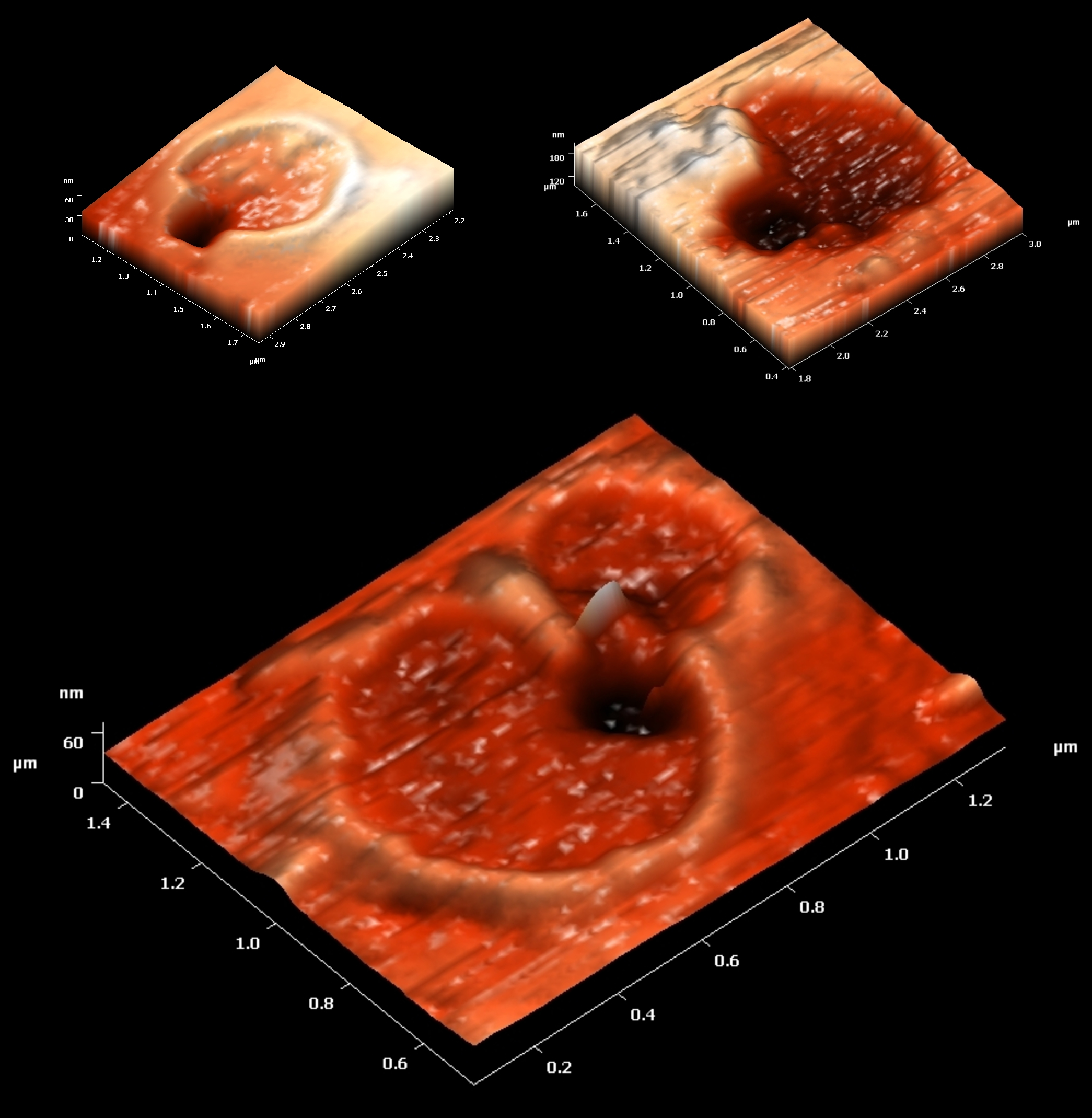

Figure 16. Examples of invaginations with additional invagination inside from donor samples (scanned in contact mode)

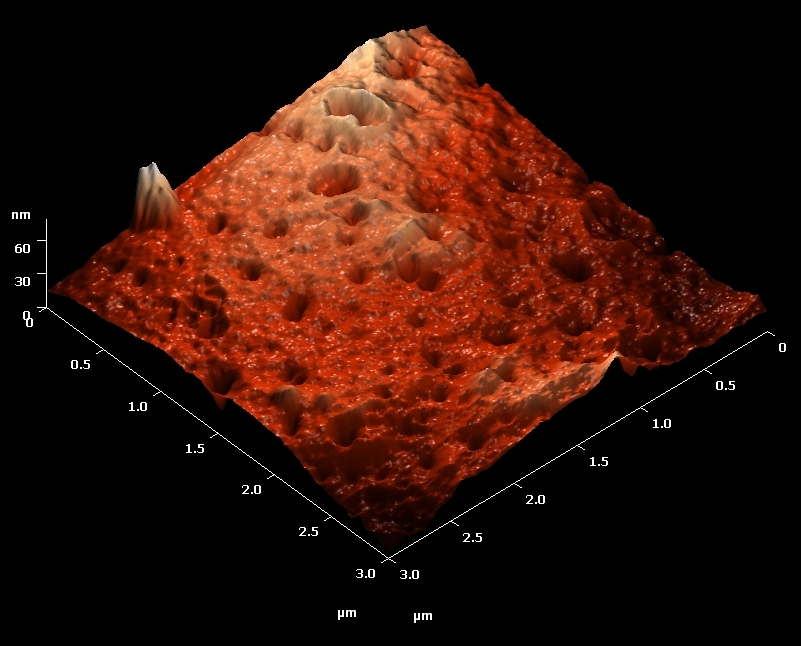

Figure 17. Examples of small invaginations (supposed endocytosis process) from AML samples (scanned in contact mode)
